# singIST: an R/Bioconductor library and Quarto dashboard for automated single-cell comparative transcriptomics analysis of disease models and humans

**DOI:** 10.64898/2026.03.03.709270

**Authors:** Aitor Moruno-Cuenca, Sergio Picart-Armada, Alexandre Perera-Lluna, Francesc Fernández-Albert

## Abstract

Preclinical disease models often diverge from human pathophysiology at single-cell resolution, complicating model selection and limiting translational value. We present `singIST`, an `R/Bioconductor` package for quantitative and explainable comparison of disease model scRNA-seq data against a human reference. For each superpathway, singIST fits an adaptive sparse multi-block PLS-DA model on human pseudobulk expression, integrated one-to-one orthology and cell type mapping, and translates model fold changes into the human expression space to compute signed recapitulation at the superpathway, cell type, and gene levels. To streamline interpretation and reporting, we provide singIST Visualizer, a companion Quarto/Shiny dashboard that loads singIST outputs and offers interactive exploration with export ready plots and tables, avoiding manual figure coding across many superpathways and models. We demonstrate the workflow. We illustrate an end-to-end workflow on an oxazolone mouse model against a human atopic dermatitis reference for two representative pathways: Dendritic Cells in regulating Th1/Th2 Development [BIOCARTA] and Cytokine-cytokine receptor interaction [KEGG]. `singIST` is distributed under the MIT License via Bioconductor, and the Visualizer is available on GitHub.

## Introduction

Translational research depends on preclinical systems that faithfully mimic human disease at molecular resolution. Bulk frameworks such as *In Silico* Treatment (IST) [1] simulate animal-model fold-changes onto human data to derive pathway recapitulation scores, and Found In Translation (FIT) [2] uses regularized regression for model-to-human alignment. However, these methods cannot resolve cell type heterogeneity, obscuring critical mechanisms in complex tissues where specific cell types drive the pathophysiology of disease. R/Bioconductor tools like CoSIA [3] and HybridExpress [4] facilitate cross-species or hybrid bulk analyses, for specific comparisons, but lack single-cell resolution and pathway-level interpretability. To address this gap, we introduce `singIST` [5], which operates on single-cell transcriptomics data, integrating human data through adaptive sparse multiblock PLS-DA (asmbPLS-DA) with one-to-one orthology and cell type mapping, to deliver interpretable recapitulation metrics between the transcriptomic changes observed in the disease model to that of a human disease at pathway, cell type, and gene levels.

## 1 Design and Implementation

`singIST` (R ≥4.6.0) is distributed via Bioconductor under the MIT License. `singIST` Visualizer is available at GitHuB. There are two main entry points for analysis and one for rendering outputs: `fitOptimal()` (and its wrapper `multiple_fitOptimal()`, `singISTrecapitulations()` (and its wrapper `multiple_singISTrecapitulations()`, `render_multiple_outputs()`. In each of the following subsections we document each in turn, including required inputs, key parameters, and returned objects. A summary of the full workflow of singIST can be found in Fig. 1.

**Fig 1.**
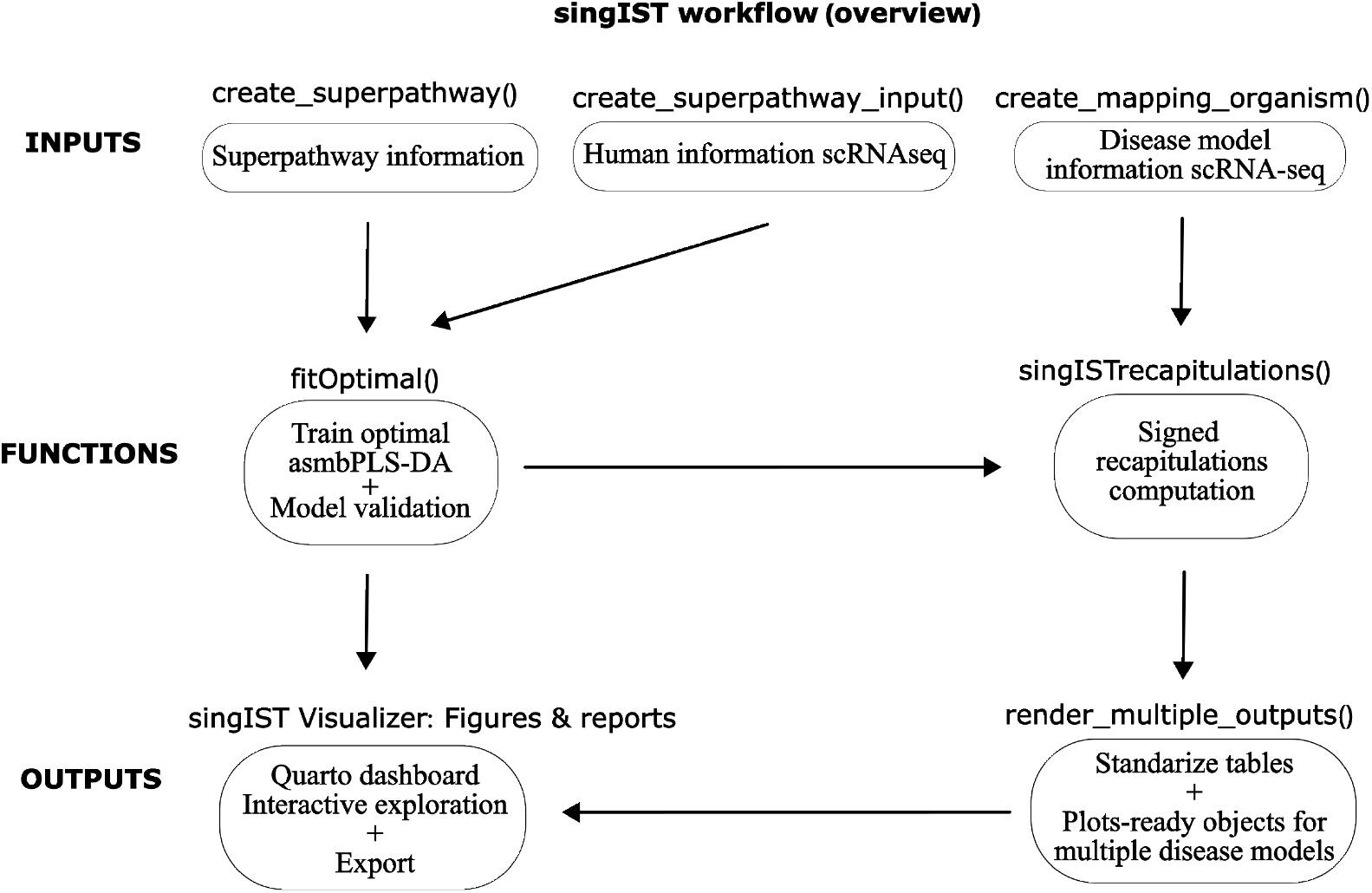
singIST workflow (overview). Input lists (superpathway definition, human reference pseudobulk data with tuning settings, and disease model scRNA-seq with cell type mapping) are created with `create_*` constructors. `fitOptimal()` trains and validates the optimal human asmbPLS-DA model, `singISTrecapitulations()` computes signed recapitulation scores, and `render_multiple_outputs()` standarizes results into plot ready tables that can be explored and exported through the Quarto-based singIST Visualizer.

### 1.1 Preparing Input Data

Before calling any of the other functions, users must:

- Define the superpathways to be analyzed. Each superpathway to be analyzed will be contained in a list created by a dedicated constructor (`create_pathway`).

~~~
my_ pathway <-create _ pathway (standard _ name =
“KEGG _ CYTOKINE _ CYTOKINE _ RECEPTOR _ INTERACTION “, dbsource =”
KEGG”, collection = “c2”, subcollection = “CP”)
~~~

The `create pathway()` constructor supports pathways from databases WIKIPATHWAYS, KEGG, PID, REACTOME and BIOCARTA. Next, the user specifies which cell types to model for that pathway and assigns a gene set to each cell type. There are two options; shared genes: use the same pathway gene set for all chosen cell types; cell type specific genes: supply a distinct gene set for each cell type. If the user chooses cell type specific genes, one must manually supply a list mapping each cell type to its own gene vector. We exemplify the case of shared genes for all cell types.

~~~
my_ superpathway <-create _ superpathway (pathway _ info = my_ pathway, celltypes = c(“T-cell”, “Dendritic ?Cells”))
*# Fetch and assign gene sets from MsigDB for each modeled cell type*
gse <-msigdb :: getMsigdb (org = “hs”, id = c(“SYM”, “EZID”), version = msigdb :: getMsigdb Versions () [1])
my_ superpathway <-setRepeatGene Sets (my_ superpathway, gse = gse)
~~~

- Make sure that the human reference data are appropriately pseudobulked and log-normalized. singIST expects a pseudobulk log-normalized matrix aggregated by sample and cell type (rows are sample *×* celltype profiles; columns are genes). singIST supports generating pseudobulk from either Seurat or SingleCellExperiment workflows (e.g., `Seurat::AggregateExpression()` for Seurat; or a dedicated SCE pseudobulk approach if working with SingleCellExperiment).

~~~
pseudobulk _ lognorm <-Seurat :: AggregateExpression (human,
   group . by = c(“celltype”, “sample ID”),
   normalization . method = “Log Normalize”) $ RNA $ data
~~~

otherwise if the object is SingleCellExperiment one can use the method

~~~
pseudobulk _ lognorm <-sing IST :: pseudobulk _ sce (human, “
   celltype”, “sample ID”)
~~~

The SingleCellExperiment object should have been previously log-normalized.

- Ensure that the disease model scRNA-seq object contains standarized metadata fields. The disease model input is a Seurat or SingleCellExperiment object and must contain:
  - `class` (experimental group membership; base vs. target)
  - `celltype_cluster` (model clusters used to map to human cell types).
  - `donor` (sample/donor identifier).

### 1.2 asmbPLS-DA fitting and validation

The first function the user will use is `fitOptimal()`, or its wrapper `multiple_fitOptimal(` if multiple superpathways are to be analysed for the same human reference. The input of `fitOptimal()` is a `superpathway_input` list and its output is a fitted-model list containing the optimal model, optimal hyperparameters and validation metrics. Before calling the function, the user must define the `superpathway`_*i*_*nputlist*.

- `First, initialize a hyperparameters list to define grid and CV design`

~~~
quantile _ comb _ table <-as. matrix (
   Rcpp Algos :: permute General (
   seq (0.05, 0.95, by = 0.50),
   m = length (my_ superpathway $ celltypes),
   repetition = TRUE
 )
)
my_ hyperparameters <-create _ hyperparameters (
   quantile _ comb _ table = quantile _ comb _table,
   outcome _ type = “binary”,
   number_ PLS = as. integer (3),
   folds _CV = as. integer (1)
)
~~~

We have specified a LOOCV approach as folds_CV = 1. We suggest to use LOOCV for small sample size. It is required to provide a number of combinations of sparsity quantiles to test for each cell type. If *b* = 1, …, *B* are the cell types modeled, 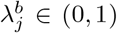 is the sparsity quantile for PLS *j*, and 0 *< w <* 1 is the step parameter along the interval (0, 1), the total number of quantiles to assess at each PLS step will be:

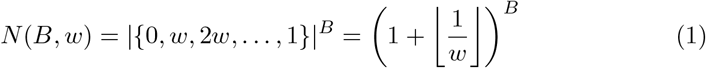

We suggest a reasonable number of combinations to be tested, although beware that the computational cost will increase exponentially as one decreases the step parameter or the number of cell types within a pathway.

- Second, initialize a superpathway input list

~~~
my_ superpathway _ input <-create _ superpathway _ input(
   superpathway _ info = my_ superpathway,
   hyperparameters _ info = my_ hyperparameters,
   pseudobulk _ lognorm = pseudobulk _ lognorm,
   sample _id = c(“AD1”, “AD2”, “HC1”, “HC2”),
   sample _ class = c(“AD”, “AD”, “HC”, “HC”),
   base _ class = “HC”,
   target_ class = “AD”
)
~~~

If the user aims to analyze only one superpathway fitOptimal() will suffice, otherwise if a list of superpathway iput lists are provided then call multiple fitOptimal(). We exemplify a fitOptimal() call with parallelization, only available when CV design is LOOCV, jackknife distribution estimation, and 100 permutations for the permutation tests of gene importance and global significance of optimal asmbPLS-DA.

~~~
*# Note that parallelization options will vary based on your OS*
library (Bioc Parallel)
Bioc Parallel :: register(Bioc Parallel :: Snow Param (workers = 2,
            exportglobals = FALSE, progressbar = TRUE),
            default = TRUE)
model <-fitOptimal(my_ superpathway _input, parallel = TRUE,
   type = “jackkniffe”, nperm = 100)
*# Disable parallelization*
Bioc Parallel :: register(Bioc Parallel :: SerialParam (),
            default = TRUE)
~~~

### 1.3 Recapitulation analysis

The second function the user will use is singISTrecapitulations(). This function receives a superpathway_fit_model_list and mapping organism list and returns a list containing: recapitulation metrics (superpathway, cell type) and gene contributions, and orthology mapping for each superpathway. This step requires:

- a fitted model list returned byfitOptimal() / multiple fitOptimal()
- a mapping_organism list created with create_mapping_organism(), which bundles the disease model information to be used

~~~
my_ mapping _ organism <-create _ mapping _ organism (
    organism = “Mus ?musculus”,
    target_ class = “ETOH”,
    base _ class = “OXA”,
    celltype _ mapping = celltype _ mapping,
    counts = object
)
~~~

The target_class and base class_slots are values contained in celltype cluster variable, and object refers to either a Seurat or SingleCellExperiment object. We call singISTrecapitulations() if only one superpathway is to be assessed against the mapping organism list, otherwise call multiple_singISTrecapitulations().

~~~
recapitulations <-sing ISTrecapitulations (model,
                                            my_ mapping _ organism
)
~~~

Internally, singISTrecapitulations() computes superpathway and cell type recapitulations, as well as gene level contributions, and returns the orthology mapping used for each superpathway.

### 1.4 Visualizing results

To facilitate interactive exploration of singIST outputs, we provide the singIST Visualizer, a Quarto-based Shiny dashboard that consumes the results of the core analysis and renders publication-quality plots and tables. This component is fully optional but recommended for rapid interrogation and sharing of findings.

The visualizer is distributed as a single Quarto document (singIST_visualizer.qmd) with Shiny runtime. After installing singIST, launch singIST Visualizer by running “singIST visualizer.qmd” file.

`It expects two inputs via the sidebar file-upload widgets:`

- Optimal models (.RData): the named list returned by fitOptimal() or multiple fitOptimal()
- Recapitulations (.RData): the output of render_multiple_outputs(recapitulations)

The dashboard loads these objects into memory and exposes three primary tabs, built with standard R packages:

- Data exploration. The first tab lets users inspect the human pseudobulk inputs with a gene-set size barplot and a block-matrix PCA (via factoextra), overlaid with boxplots and individual sample IDs (DT, ggplot2) for any selected gene-and-cell type combination.
- Optimal model. The second tab focuses on model performance and feature importance. A side-by-side boxplot (via ggplot2) presents the distribution of superpathway scores for the base versus target classes, annotated with BH-adjusted p-values derived from the CV scheme chosen in fitOptimal(). Adjancent radar chart, generated with ggradar, summarize Cell Importance Projections (CIP) across cell type blocks. Below, an interactive jitter plot, built with plotly, displays the top N genes --ranked by Gene Importance Projection (GIP magnitude and −log10 adjusted p-value --for any selected cell type, enabling users to identify the most predictive markers. Two patterns can emerge in this jitter plot; cluttered genes, it is a group of genes that drive the prediction; scattered genes, among the statistically significant genes there are genes with much higher predictive power. A searchable DataTable lists gene names, GIP values, direction of effect, and adjusted p-values, with copy/CVS/Excel export buttons.
- Recapitulations. The final tab presents the core outcome of singISTrecapitulations(). A heatmap of fractional superpathway recapitulation 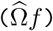 is drawn with ComplexHeatmap, with each tile colored on a blue-white-green gradient and annotated with percentage values. Below, one-to-one orthology percentages for each cell type gene set are shown as grouped bar charts via ggplot2, facilitating assessment of cross-species mapping fidelity. A faceted tile plot then displays cell type recapitulation scores 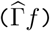 for each pathway and model, with labels indicating percentages. Finally, a composite ComplexHeatmap juxtaposes gene-level contributions 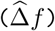 and model fold-changes 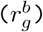 for the top drivers genes per cell type: contributions are colored on a red-white-blue scale, while fold-chang use a separate palette and border annotations mark statistical significance. An accompanying DataTable allows users to sort and filter genes by contribution, fold-change, orthology status, and adjusted p-value.

By combining all these three interactive panels, the singIST Visualizer transforms the raw analysis outputs into an intuitive user experience, enabling rapid hypothesis generation and automated reporting of single-cel model evaluation.

## 2 Results

We showcase singIST on the deposited test datasets distributed with the package, consisting of:

- a human scRNA-seq reference summarized as cell type pseudobulk log-normalized expression profiles for healthy control versus lesiona skin of atopic dermatitis patients
- a disease model scRNA-seq (Seurat) annotated with donor, condition (OXA vs. ETOH), and model specific cell type clusters.

We define two examples of superpathways from curated MsigDB collections and fit asmbPLS-DA atopic dermatitis human reference model with CV and permutation based validation, using the parameters provided in the vignette. The fitted reference models are then used to quantify disease model to human recapitulation at superpathway, cell type, and gene level, and all outputs are rendered into standarized objects for downstream plotting and interactive exploration in singIST Visualizer with Quarto.

On the “Dendritic Cells in Th1/Th2 Development [BIOCARTA]” superpathway the oxazolone mouse model shows moderate superpathway recapitulation (52.4%), whereas “Cytokine-cytokine receptor interaction [KEGG]” almost null recapitulation (8.5%) (Fig. 2A).

**Fig 2.**
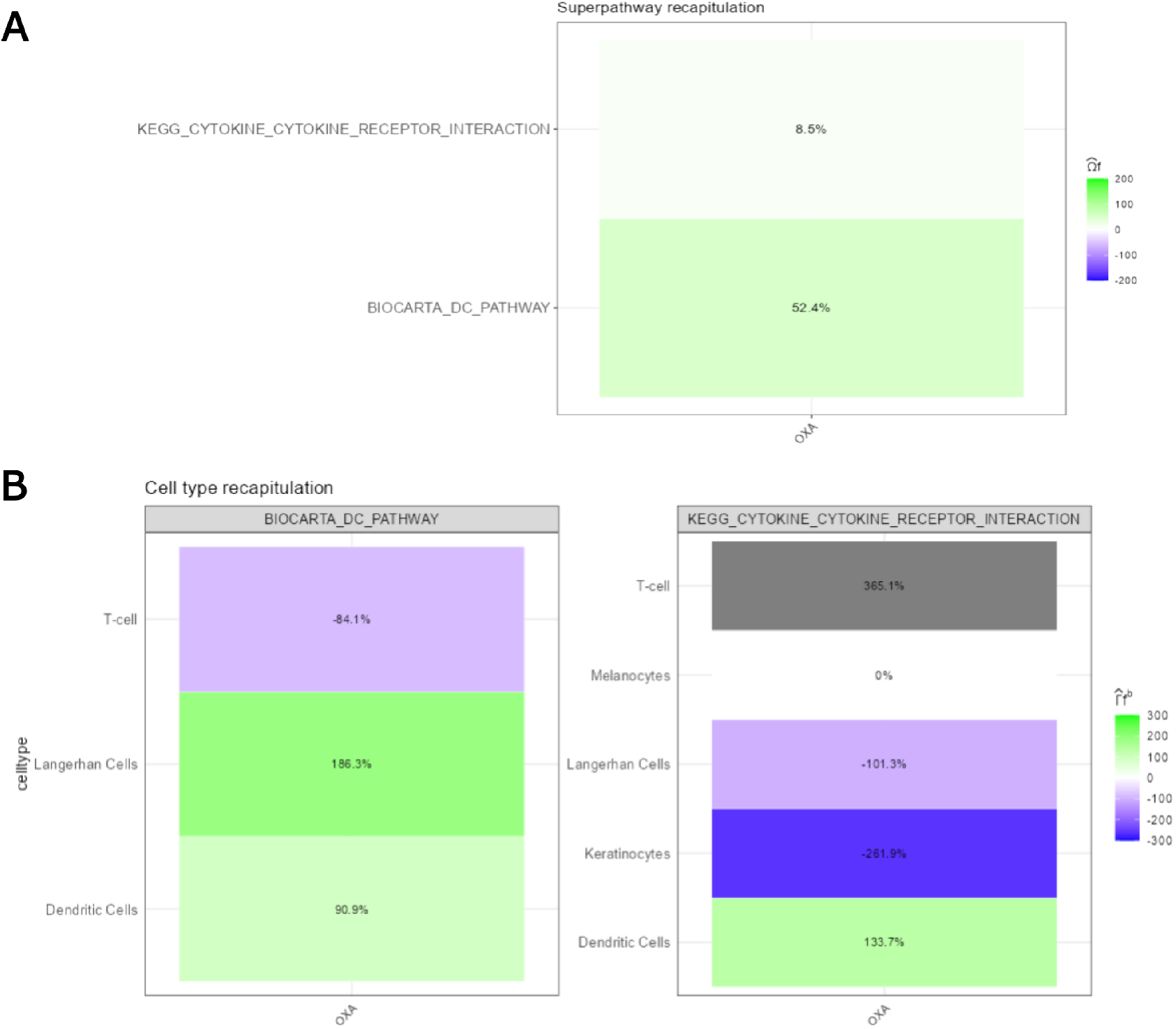
Oxazolone mouse model recapitulations against Atopic Dermatitis human reference. A) Superpathway recapitulation 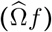. Signed fractional recapitulation at the superpathway level for two superpathways in the oxazolone (OXA) model. Values quantify how closely disease model induced changes align with the direction and magnitude of the human reference signal; negative values indicate opposite direction shifts and values greater than 100% indicate larger magnitude shift. **B) Cell type recapitulation** 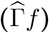 Cell type signed fractional recapitulation for the same two superpathways. The heatmaps highlight which cell types drive superpathway level agreement or disagreement, and reveal opposing recapitulations that can partially cancel at the superpathway level.

Because superpathway level recapitulations can mask heterogeneous cell type contributions, we check the cell type recapitulations in Fig. 2B. For the BIOCARTA pathway, recapitulation is driven by strong aligned shifts in Langerhans Cells (186.3%) and Dendritic Cells (90.9%), while T-cell shows an opposite direction contribution (−84.1%), indicating that partial superpathway recapitulation can arise from mixed cell type shifts. The KEGG pathway, shows strong positive recapitulations for T-cell and Dendritic Cells (365.1%; 133.7%), counterbalanced by negative recapitulations in Keratinocytes (−261.9%) and Langerhans Cells (−101.3%).

The vignette of these results can be found in Supplementary Material S1.

## 3 Availability and Future Directions

singIST (R ≥ 4.6.0) is distributed via Bioconductor under the MIT License. Package can be installed via BiocManager::install(“singIST”).

The singIST Visualizer can be installed via remotes::install_github(“DataScienceRD-Almirall/singIST-visualizer”)

## Supporting information

S1. File. Supplementary Material S1 - Results Vignette

This supplementary material is an .Rmd file with the singIST workflow generated for the results section.

## Acknowledgments

The funders had no role in study design, data collection and analysis, decision to publish, or preparation of the manuscript. This work was supported by the Spanish Ministry of Economy and Competitiveness (www.mineco.gob.es) PID2021-122952OB-I00 and DPI2017-89827-R, and Doctorats Industrials Agència de Gestió d’Ajuts i Universitaris i de Recerca (AGAUR) 2023 DI 00053.

