## Supplementary material for "singIST: an R/Bioconductor library and Quarto dashboard for automated single-cell comparative transcriptomics analysis of disease models and humans": S1 File: SP1_singISTvignette.Rmd

````
---
title: "singIST Vignette"
author: "Aitor Moruno-Cuenca"
output: BiocStyle::html_document
vignette: > 
  %\VignetteIndexEntry{singIST Vignette}
  %\VignetteEngine{knitr::rmarkdown}
  %\VignetteEncoding{UTF-8}
---

# Abstract of singIST as a Bioconductor library
Comparative transcriptomics aims to identify common and differing patterns
(organisms, experimental conditions, tissues, and so on) between a set of
conditions by using transcriptomics data The methodology to tackle the problem
will vary greatly depending on which patterns we aim to compare, the amount and
biological nature of them and which omics resolution. **singIST** is a
comparative transcriptomics method that aims to compare a disease model against
a human disease of interest. In drug discovery and pharmaceutical early
development choosing a disease model becomes a fundamental endavour. For this
reason, we present **singIST** as a method that compares the single-cell
transcriptomics profile of a disease model, either In Vivo or In Vitro models,
against a human disease that such model should mimick.

Other Bioconductor packages for comparative transcriptomics are available.
One of such is **HybridExpress** which aims to compare the bulk transcriptomics
(RNA-seq) profile for hybrids relative to their progenitor species. **CoSIA**
is a Bioconductor package that enables organism-to-organism comparative bulk
transcriptomics (RNAseq) across tissues. However, none of these packages are
direct comparators to **singIST* since they are not conceived for single-cell
comparative transcriptomics but for bulk transcriptomics instead.


# Introduction

This Vignette demonstrates how to use the singIST package. singIST quantifies
the direction and mangitude of single-cell transcriptomic changes between a
human reference condition and a disease model at pathway, cell type, and gene
resolution. Throughout this vignette we will work with the following data:

- "cytokine_counts.rda" Contains Human scRNAseq pseudobulk LogNormalized
count matrix. The pseudobulk is aggregated by sample and cell type and contains
all genes belonging to the pathway Cytokine-Cytokine receptor interaction [KEGG]
- "DC_counts.rda" Contains Human scRNAseq pseudobulk LogNormalized
count matrix. The pseudobulk is aggregated by sample and cell type and contains
all genes belonging to the pathway Dendritic Cells in regulating Th1/Th2
Development [BIOCARTA]
- "OXA.rda" 

```{r}
set.seed(003)
library(singIST)
```

# Inputs of singIST

The method is defined around three inputs:

- **Superpathways**: pathway gene sets structured by cell type. 
- **Human scRNA-seq reference (pseudobulk)**: transformed into cell-type
specific pseudobulk, restricted to the genes of the superpathway.
- **Disease model scRNA-seq**: a Seurat or SingleCellExperiment object
containing scRNA-seq readouts of the disease model to assess.

In the following subsections we explicitly construct these inputs. The rest of
the vignette uses these inputs to: (1) learn a human reference model using 
adaptive sparse multi-block PLS-DA, and then (2) evaluate how human-like the
disease model changes are, in the learned human reference space.

## Superpathways

singIST does not operate on a pathway gene set alone. Instead, it defines a
superpathway by three things:

\begin{itemize}
\item A curated **pathway** (e.g., an MSigDB pathway identifier).
\item A set of **cell types** of interest that will be modeled explicitly.
\item A **gene set per cell type**, where each cell type is associated with a 
vector of genes that might or might not be the same
\end{itemize}

In many practical cases, we use the same pathway gene set for every cell type. 
However, singIST also supports **cell-type specific gene sets**. Below we create
two example of superpathways used later in this vignette.

First, because singIST relies on MSigDB curated pathways, we load the MSigDB
gene set collection once and reuse it as necessary. We assume the reference
to compare against is human `org = "hs"` and we extract both ENTREZID
and HGNC gene symbols `id = c("SYM", "EZID")`.

```{r, echo = TRUE, message = FALSE, warning = FALSE}
gse <- msigdb::getMsigdb(org = "hs", id = c("SYM", "EZID"),
                         version = msigdb::getMsigdbVersions()[1])
```

### Example 1: Superpathway with the same pathway gene set for each cell type

In this first example, we define a pathway using its MSigDB metadata, choose the
cell types we want to model, and then assign the same pathway gene set to each
cell type gene set. This is the simplest and most common setup for a 
superpathway.

```{r, echo = TRUE, message = FALSE, warning = FALSE}
# Pathway definition (curated gene set identifier)
dc_pathway <- create_pathway(standard_name = "BIOCARTA_DC_PATHWAY",
                             dbsource = "BIOCARTA",
                             collection = "c2",
                             subcollection = "CP")
# Superpathway = pathway + modeled cell types + per cell type gene sets
dc_superpathway <- create_superpathway(pathway_info = dc_pathway,
                                       celltypes = c("Dendritic Cells",
                                                     "Langerhan Cells",
                                                     "T-cell"),
                                       list())
# Assign gene sets per cell type (retrieve gene set from MSigDB)
dc_superpathway <- setGeneSetsCelltype(dc_superpathway, gse = gse)
```

### Example 2: Superpathway with different gene sets than the retrieved (optional)

In some applications, you may want different cell types to use a subset or 
different thereof. singIST supports this by allowing the user to directly
provide a list of gene vectors, one per cell type. 

Here we demonstrate this capability by taking the curated pathway gene set and 
creating different subsets for the cell type. (This is purely illustrative;
in real analyses these subsets would be defined biologically rather than 
randomly sampled.)

```{r, echo = TRUE, message = FALSE, warning = FALSE}
# Pathway definition (curated gene set identifier)
cytokine_pathway <- create_pathway(
    standard_name = "KEGG_CYTOKINE_CYTOKINE_RECEPTOR_INTERACTION",
    dbsource = "KEGG",
    collection = "c2",
    subcollection = "CP"
)
# Superpathway = pathway + modeled cell types + per cell type gene sets
cytokine_superpathway <- create_superpathway(
    pathway_info = cytokine_pathway,
    celltypes = c("Dendritic Cells", "Keratinocytes", "Langerhan Cells",
                  "Melanocytes", "T-cell"),
    list()
)
# Pull the curated gene set (retrieve gene set from MSigDB)
gene_set <- pullGeneSet(cytokine_pathway, gse = gse)
sets <- sample(gene_set, 100)
# Provide a gene set per cell type
cytokine_superpathway$gene_sets_celltype <- list(
    "Dendritic Cells" = sets,
    "Keratinocytes" = sets,
    "Langerhan Cells" = sets,
    "Melanocytes" = sets,
    "T-cell" = sets
)
```

## Human scRNA-seq

The human reference data are used to train an adaptive sparse multi-block PLS
on a superpathway. In this subsection we start from the human scRNA-seq that
have already been aggregated into cell-type pseudobulk log-normalize expression.
For instance, the pseudobulk can be performed with 
`Seurat::AggregateExpression()`. Concretely the pseudobulk matrix:

- Each row corresponds to a pseudobulk profile for a cell type within a sample
(e.g., T-cell_AD1).
- Columns correspond to genes (typically HGNC symbols for human training).
- Values correspond to log-normalized pseudobulk expression.

In addition to the pseudobulk matrix, singIST requires sample level labels
defining a base class and a target class (e.g., HC vs AD). The target class
is the human experimental group that the disease model such mimic precisely, 
while the base class is the human experimental group that such differentiate 
from the target class.

The training procedure includes a tuning step, so the human input is not only 
the pseudobulk matrix and sample labels - it also includes the hyperparameter
search space and the cross-validation design the user wants. In practice,
we tune:

- Block-specific sparsity via a grid of quantile thresholds (one quantile
per modeled cell type). Intuitively, higher quantiles enforce stronger sparsity
(fewer genes retained) in that block.
- Model complexity via `number_PLS` (maximum number of latent components
to test).
- Validation scheme via `folds_CV` and `repetition_CV`.

We store the former settings in a `hyperparameters` list and then combine
all the information into the `superpathway_input` list, which is the main
training input consumed by the training procedure `fitOptimal()/
multiple_fitOptimal()`.

### Example 1: Dendritic Cells in regulating Th1/Th2 Development

```{r, echo = TRUE, message = FALSE, warning = FALSE}
# Load the pseudobulk matrix
file_dc <- system.file("extdata", "DC_counts.rda", package = "singIST")
obj_dc <- load(file_dc)
counts_dc <- get(obj_dc[1])
# Remove the cell types that we are not going to model
counts_dc <- counts_dc[!(sub("_.*$", "", rownames(counts_dc)) %in% 
                        c("MastC-other", "Keratinocytes", "Melanocytes")), ]
```

Next, we define the sample identifiers and class labels.

```{r, echo = TRUE, message = FALSE, warning = FALSE}
# Load the pseudobulk matrix
sample_id_dc <- c("AD1", "AD13", "AD2", "AD3","AD4", "HC1-2", "HC3", "HC4",
                  "HC5")
sample_class_dc <- c(rep("AD", 5), rep("HC", 4))
base_class_dc <- "HC"
target_class_dc <- "AD"
```

Now we define the hyperparameters.

```{r, echo = TRUE, message = FALSE, warning = FALSE}
# Initialize hyperparameter object 
# Choose the array of quantile combinations that will be tested in the Cross
# Validation. Ideally one should choose a wide range of quantiles to test,
# as a narrow selection might not find the globally optimal model. The more
# quantiles one chooses the more computation time it will take, for the sake
# of simplicity we choose a small amount of quantiles to test
quantile_comb_table_dc <- as.matrix(
    RcppAlgos::permuteGeneral(seq(0.1, 0.95, by = 0.3),
                                m = length(dc_superpathway$celltypes), 
                                TRUE))
# Store tuning + CV design
dc_hyperparameters <- create_hyperparameters(
    quantile_comb_table = quantile_comb_table_dc,
    outcome_type = "binary",
    number_PLS = as.integer(3),
    folds_CV = as.integer(1), #LOOCV
    repetition_CV = as.integer(1)
)
```

Finally, we combine all the information into a superpathway_input list.

```{r, echo = TRUE, message = FALSE, warning = FALSE}
dc_superpathway_input <- create_superpathway_input(
    superpathway_info = dc_superpathway,
    hyperparameters_info = dc_hyperparameters,
    pseudobulk_lognorm = counts_dc,
    sample_id = sample_id_dc,
    sample_class = sample_class_dc,
    base_class = base_class_dc,
    target_class = target_class_dc
)
```


### Example 2: Cytokine-Cytokine receptor interaction superpathway

```{r, echo = TRUE, message = FALSE, warning = FALSE}
# Load the pseudobulk matrix
file_cytokine <- system.file("extdata", "cytokine_counts.rda", package = "singIST")
obj_cytokine <- load(file_cytokine)
counts_cytokine <- get(obj_cytokine[1])
# Remove the cell types that we are not going to model
counts_cytokine <- counts_cytokine[!(sub("_.*$", "", rownames(counts_cytokine)) 
                                     %in% 
                                 c("MastC-other")), ]
```


Next, we define the sample identifiers and class labels.

```{r, echo = TRUE, message = FALSE, warning = FALSE}
# Load the pseudobulk matrix
sample_id_cytokine <- c("AD1", "AD13", "AD2", "AD3","AD4", "HC1-2", "HC3",
                        "HC4", "HC5")
sample_class_cytokine <- c(rep("AD", 5), rep("HC", 4))
base_class_cytokine <- "HC"
target_class_cytokine <- "AD"
```

Now we define the hyperparameters.

```{r, echo = TRUE, message = FALSE, warning = FALSE}
# Initialize hyperparameter object 
quantile_comb_table <- as.matrix(
    RcppAlgos::permuteGeneral(seq(0.05, 0.95, by = 0.2),
                                m = length(cytokine_superpathway$celltypes), 
                                TRUE))
# Store tuning + CV design
cytokine_hyperparameters <- create_hyperparameters(
                                    quantile_comb_table = quantile_comb_table,
                                    outcome_type = "binary",
                                    number_PLS = as.integer(2),
                                    folds_CV = as.integer(5),
                                    repetition_CV = as.integer(5))
```

Finally, we combine all the information into a superpathway_input list.

```{r, echo = TRUE, message = FALSE, warning = FALSE}
cytokine_superpathway_input <- create_superpathway_input(
    superpathway_info = cytokine_superpathway,
    hyperparameters_info = cytokine_hyperparameters,
    pseudobulk_lognorm = counts_cytokine,
    sample_id = sample_id_cytokine,
    sample_class = sample_class_cytokine,
    base_class = base_class_cytokine,
    target_class = target_class_cytokine)
```


## Disease model scRNA-seq

The disease model input is a single-cell object (either a `Seurat` or
`SingleCellExperiment`) containing scRNA-seq measurements from the model
system we want to assess. Unlike the human input (which is provided as 
pseudobulk matrices), the disease model is kept at single-cell resolution and is
summarized internally by singIST when computing recapitulation scores.

To use a disease model dataset, singIST needs:

- A class label indicating the base and target experimental groups
in the model (e.g. ETOH vs OXA).
- A cell-type cluster label used to map model cell populations to the human
cell types defined in the superpathway.
- A donor / sample identifier.

Finally, we must provide a cell type mapping from th ehuman superpathway cell 
types (names used in `celltypes = ...`) to the disease model clusters.
A human cell type can map to one or more model clusters. A human cell type
can map to one or more model clusters; it is also valid to provide an empty
mapping (meaning that block will not contribute for that model).

In the following example we load the oxazolone mouse dataset shipped in the
package, standarize metadata column names to those expected by singIST,
define the mapping, and construct a `mapping organism` list via
`create_mapping_organism()`.

### Example: Oxazolone mouse model of Atopic Dermatitis

```{r, echo = TRUE, message = FALSE, warning = FALSE}
file_oxa <- system.file("extdata", "OXA.rda", package = "singIST")
obj_oxa <- load(file_oxa)
oxazolone <- get(obj_oxa[1])
```
Next, we ensure the object contains the metadata fields required by singIST.
The package example dataset already contains the necessary information,
but we rename the relevant columns to the standarized names used downstream:
`class`, `celltype_cluster`, and `donor`.

```{r, echo = TRUE, message = FALSE, warning = FALSE}
# The example object included in singIST uses these column positions
colnames(slot(oxazolone, "meta.data"))[c(5,11,1)] <- c("class",
                                                       "celltype_cluster",
                                                       "donor")
# If the object is a SingleCellExperiment, metadata live in colData()
```

Now, we define the human to model cell type mapping. The names of the list
must match the human cell types used in our superpathways. Values correspond
to one or more model clusters present in `celltype_cluster`.

```{r, echo = TRUE, message = FALSE, warning = FALSE}
celltype_mapping <- list(
                    "Dendritic Cells" = c("cDC2", "cDC1", "migratory DCs"),
                    "Keratinocytes" = c("Keratinocytes"),
                    "Langerhan Cells" = c("LC"), 
                    "Melanocytes" = c(),
                    "T-cell" = c("DETC", "T"))
```

Finally, we combine the model object and mapping into a
`mapping_organism` list. This list specifies the model organism we 
are working with, and which class labels correspond to the base and
target conditions.

```{r, echo = TRUE, message = FALSE, warning = FALSE}
oxa_org <- create_mapping_organism(organism = "Mus musculus", 
                target_class = "OXA", base_class = "ETOH",
                celltype_mapping = list(
                    "Dendritic Cells" = c("cDC2", "cDC1", "migratory DCs"),
                    "Keratinocytes" = c("Keratinocytes"),
                    "Langerhan Cells" = c("LC"), 
                    "Melanocytes" = c(),
                    "T-cell" = c("DETC", "T")),
                counts = oxazolone)
```

# Fit Optimal asmbPLS-DA and compute validation metrics for the optimal model

At this stage we have constructed two key training inputs:
`dc_superpathway_input` and `cytokine_superpathway_input`. The
next step is to train an adaptive sparse multi-block PLS-DA model (asmbPLS-DA)
for each superpathway, while selecting an optimal configuration (sparsity
per cell type block and number of PLS components) under the cross-validation
design specified in the corresponding `hyperparameters` object.

singIST provides two entry points to do it:

- `fitOptimal()` fits one `superpathway_input` list.
- `multiple_fitOptimal()` is a convenience wrapper that fits multiple
superpathways in one call. 

If the CV design is a leave-one-out CV (LOOCV), the functions provide with a
parallelization option to make the computations faster via using 
`parallel = TRUE` and setting the desired paralellization scheme with 
`BiocParallel::register()`.

In addition to fitting the optimal model, singIST can compute model validation
metrics, including permutation based global significance test
(`npermut = 100`) and resampling permutation tests (`type`, e.g.,
jackknife or subsampling). The number of permutation for the tests can be chosen
as desired.

## Example: fitting cytokine-cytokine receptor interaction and dendritic cells
## in regulating 

If the human pseudobulk input contains missing values (e.g., a cell type
block absent for some samples), `fitOptimal() / multiple_fitOptimal()`
automatically imputes them during model fitting. Further details of the imputation
procedure are provided in the Appendix.

```{r, echo = TRUE, message = FALSE, warning = FALSE}
# Register a parallel backend (SnowParam works on Windows/macOS/Linux);
# on Linux you may prefer MulticoreParam.
BiocParallel::register(BiocParallel::SnowParam(workers = 2,
                        exportglobals = FALSE, progressbar = FALSE),
                        default = TRUE) 
# Combine all human training inputs into a list
superpathways <- list(dc_superpathway_input,
                      cytokine_superpathway_input)
# Fit Optimal asmbPLS-DA models + compute validation metrics
# parallel: logical vector indicating whether the CV design should be parallel
# npermut: number of permutations used for permutation based tests
# type : validation strategy (jackknife or subsampling)
# nsubsampling: number of subsampling repeats (only used when type includes
# subsampling).
superpathways_fit <- multiple_fitOptimal(
    superpathways,
    parallel = c(TRUE, FALSE), 
    npermut = c(100),
    type = c("jackknife","subsampling"), 
    nsubsampling = c(NULL, 5)
    )
# Restore serial backend (good practice)
BiocParallel::register(BiocParallel::SerialParam(), default = TRUE) 
```


# Compute recapitulation metrics

After fitting the optimal human reference models (`superpathway_fit`),
the final step is to quantify how well a disease model recapitulates the
human target vs base signal for each superpathway. singIST does this by
applying the disease model changes to the human base class reference
and then evaluating how "target-like" the samples become. The resulting
recapitulation outputs can be summarized at pathway, cell type, and gene
resolution.

There are two functions to do this:

- `singISTrecapitulations()` computes recapitulations for one fitted
superpathway model.
- `multiple_singISTrecapitulations()` is a wrapper that computes 
recapitulations for multiple fitted models at once.

Below we compute recapitulations for both fitted superpathways against the
oxazolone mapping organism `oxa_org`, and then render the results
into a plotting friendly output

## Example: Evaluating Oxazolone mouse model of Atopic Dermatitis for Dendritic
## Cells and cytokine superpathways

By default, the function assumes the organism for which the asmbPLS-DA model
was trained is "hsapiens", it also assumes that the gene annotation for the
mapped organism is "external_gene_name". These parameters are by default
but other cases can also be considered by specifying the parameters
`from_to = "hsapiens"` and
`object_gene_identifiers = "external_gene_name"`.

```{r, echo = TRUE, message = FALSE, warning = FALSE}
# Compute singIST recapitulations for all fitted superpathways
recapitulations_multiple <- multiple_singISTrecapitulations(oxa_org,
                                                            superpathways_fit)
```

In many workflows you will want the outputs in a standarized format that is
convenient for downstream plotting and reporting. The helper function
`render_multiple_outputs()` reshapes the recapitulation objects
accordingly. It makes a list that is ready-to-plot. 

```{r, echo = TRUE, message = FALSE, warning = FALSE}
final_output <- render_multiple_outputs(objects= list(recapitulations_multiple))
```

# Appendix: Missing value handling in human pseudobulk matrices

During CV, singIST may encounter missing entries in the human pseudobulk
predictor (e.g., entire cell type blocks missing for some samples). Rather
than discarding those samples, singIST imputes missing values inside each CV
split in a way that preserves the multiblock structure and avoids information
leakage. This procedure is performed automatically by `fitOptimal()`
and `multiple_fitOptimal()` and the user does not provide any input on
this, this section is merely for informative purposes.

## Problem setup

Assume we have a multi-block dataset for which we aim to impute an asmbPLS-DA,
$D = \{\left(x_i, y_i\right) \}_{i=1}^n$ where $x_i \in \mathbb{R}^p, y_i \in \{0,1\}$.
We perform a K-fold CV, thus splitting the dataset into $K$ folds; $\mathcal{T}_k$
training indices, $\mathcal{V}_k$ validation indices, $k = 1, \dots, K$. 
Then we have a classic train-test dataset
$\left(X_{train}^{(k)}, Y_{train}^{(k)}  \right) \in \{(x_i, y_i): i \in \mathcal{T}_k \}$
and $\left(X_{val}^{(k)}, Y_{val}^{(k)}  \right) \in \{(x_i, y_i): i \in \mathcal{V}_k \}$

The problem lies in that $X$ matrices we might have missing values.

## Step 1: Impute the training fold using multiblock MFA.

We first impute $X_{train}^{(k)}$ using Multple Factor Analysis (MFA) imputation
(via `missMDA::imputeMFA()`), which explicitly accounts for the 
multiblock structure (blocks correspond to modeled cell types). This produces
an imputed training matrix $\widetilde{X}_{train}^{(k)}$.

## Step 2: Impute the validation fold by projection (no re-fitting).

To avoid data leakage, we do not refit the MFA model on the validation data.
Instead, we compute MFA loadings and means from the imputed training fold and
project each validation sample using its observed features only.

Let $\mu \in \mathbb{R}^p$ be the vector of variable means and
$P \in \mathbb{R}^{p \times ncp}$ the matrix of MFA loadings estimated from
$\widetilde{X}_{train}^{(k)}$. For each validation sample
$x_i \in X_{val}^{(k)}$ define the set of observed feature indices:

$$O_i = \{j: (x_i)_j \neq NA\}, \quad |O_i| \geq 1$$

We estimate the latent scores $t_i$ by solving a least-squares problem 
restricted to the observed features:

$$t_i = \arg \min_{u_i \in \mathbb{R}^{ncp}} ||x_{i, O_i}-\left(\mu_{O_i}+P_{O_i}u_i \right)||_2^2 = \left(P_{O_i}^TP_{O_i}\right)^{-1} P_{O_i}^T \left(x_{i,O_i}-\mu_{O_i} \right)$$

We then reconstruct the feature full vector by:

$$\widehat{x}_{ij} = \mu_{j} + P_{j\cdot} t_i$$

The imputation validation entries are defined as:

$$
\begin{align}
\widehat{x}_{ij}  = \begin{cases}
x_{ij}, & j \in O_i\\
\widetilde{x}_{ij}, & j \notin O_i
\end{cases} \quad i \in \mathcal{V}_k
\end{align}
$$

This preserves the same latent space used for training-fold imputation and
prevents leakage from validation into the imputation model. Imputation is
intended for **partial missingness** (e.g., one missing block while other
blocks are observed). If missingness is extensive (e.g., many samples missing
most blocks), model stability and interpretability may degrade; in that case
consider restricting the set of modeled cell types and/or increasing sample
size.

# Session Info

```{r}
utils::sessionInfo()
```
````
